# Trans-splicing of the *C. elegans let-7* primary transcript developmentally regulates *let-7* microRNA biogenesis and *let-7* family microRNA activity

**DOI:** 10.1101/400077

**Authors:** Charles Nelson, Victor Ambros

## Abstract

*let-7* is a microRNA whose sequence and roles as a regulator of developmental progression are conserved throughout bilaterians. In most systems, transcription of the *let-7* locus occurs relatively early in development, whilst processing of *let-7* primary transcript into mature microRNA arises later and is associated with cellular differentiation. In *C. elegans* and other animals, the RNA binding protein LIN-28 post-transcriptionally inhibits *let-7* biogenesis at early developmental stages. The mechanisms by which LIN-28 and other factors developmentally regulate *let-7* biogenesis are not fully understood. Nor is it understood how the developmental regulation of *let-7* might influence the expression or activities of other microRNAs of the same seed family. Here we show that in *C. elegans*, the primary *let-7* transcript (*pri-let-7*) is trans-spliced to SL1 splice leader at a position downstream of the *let-7* precursor stem-loop, producing a short, polyadenylated downstream mRNA. The trans-splicing event negatively impacts the biogenesis of mature *let-7* microRNA in *cis*, likely by destabilizing the upstream *pri-let-7* fragment. Moreover, the trans-spliced downstream mRNA contains complimentary sequences to multiple members of the *let-7* seed family (*let-7fam*), and thereby serves as a sponge to negatively regulate *let-7fam* function in *trans*. Thus, this study provides evidence for a mechanism by which splicing of a microRNA primary transcript can negatively regulate said microRNA in *cis* as well as other microRNAs in *trans*.

**HIGHLIGHTS:**

- The *let-7* primary transcript is trans-spliced to produce an RNA that functions as a sponge that negatively regulates the *let-7-family* microRNAs.
- Trans-splicing of this RNA negatively impacts *let-7* microRNA biogenesis.
- LIN-28 regulates this trans-splicing event

## INTRODUCTION

MicroRNAs are endogenous ∼22 nucleotide RNAs that are enzymatically processed from longer primary transcripts, and that repress protein expression through imperfect base pairing with their target mRNAs. Nucleotides 2-8 of the microRNA, known as the seed, instigate target recognition through essentially complete complementarity, whereas base pairing via the non-seed nucleotides (9-22 of the microRNA) is less constrained than is seed pairing (He and Hannon, 2004). microRNAs that contain an identical seed sequence but differ in their non-seed nucleotides are classified together as a “family” based on their presumed evolutionary relatedness, and their potential to act in combination on the same targets (Bartel, 2009, Ambros and Ruvkun, 2018).

The *let-7* gene was initially identified in a screen for developmental defects in *Caenorhabditis elegans* (Meneely and Herman, 1979), and later found to encode a microRNA that promotes the differentiation of cellular fates (Reinhart et al., 2000). Orthologs of the *C. elegans let-7* microRNA are easily identified across animal phyla because of the near perfect conservation of the entire 22 nt sequence (Pasquinelli et al., 2000). In many species, *let-7* paralogs encode additional family members, including *mir-48*, *mir-84* and *mir-241* in *C. elegans,* that differ from *let-7* in some of their non-seed nucleotides. In *C. elegans*, *let-7fam* microRNAs function semi-redundantly to regulate stage-specific larval cell fate transitions, with *miR-48, mir-84* and *mir-241* primarily promoting the L2-to-L3 transition and *let-7* primarily promoting the L4-to-adult transition (Reinhart et al., 2000, Abbott et al., 2005).

In *C. elegans*, two major primary transcripts of *let-7* are produced, pri*-let-7 A* and *pri-let-7 B* of 1731 and 890 nucleotides, respectively. The 5’ end of these transcripts can be further processed by SL1 trans-splicing to produce the 728 nucleotide *SL1-pri-let-7* (Bracht et al., 2004), (Fig. 1A). All three of these *pri-let-7* transcripts contain the *let-7* precursor hairpin plus additional downstream sequences, including an element with complementarity to the *let-7fam* seed sequence that has been shown to associate in *vivo* with the Argonaut protein, ALG-1 (Zisoulis et al., 2012).

**Figure 1.**
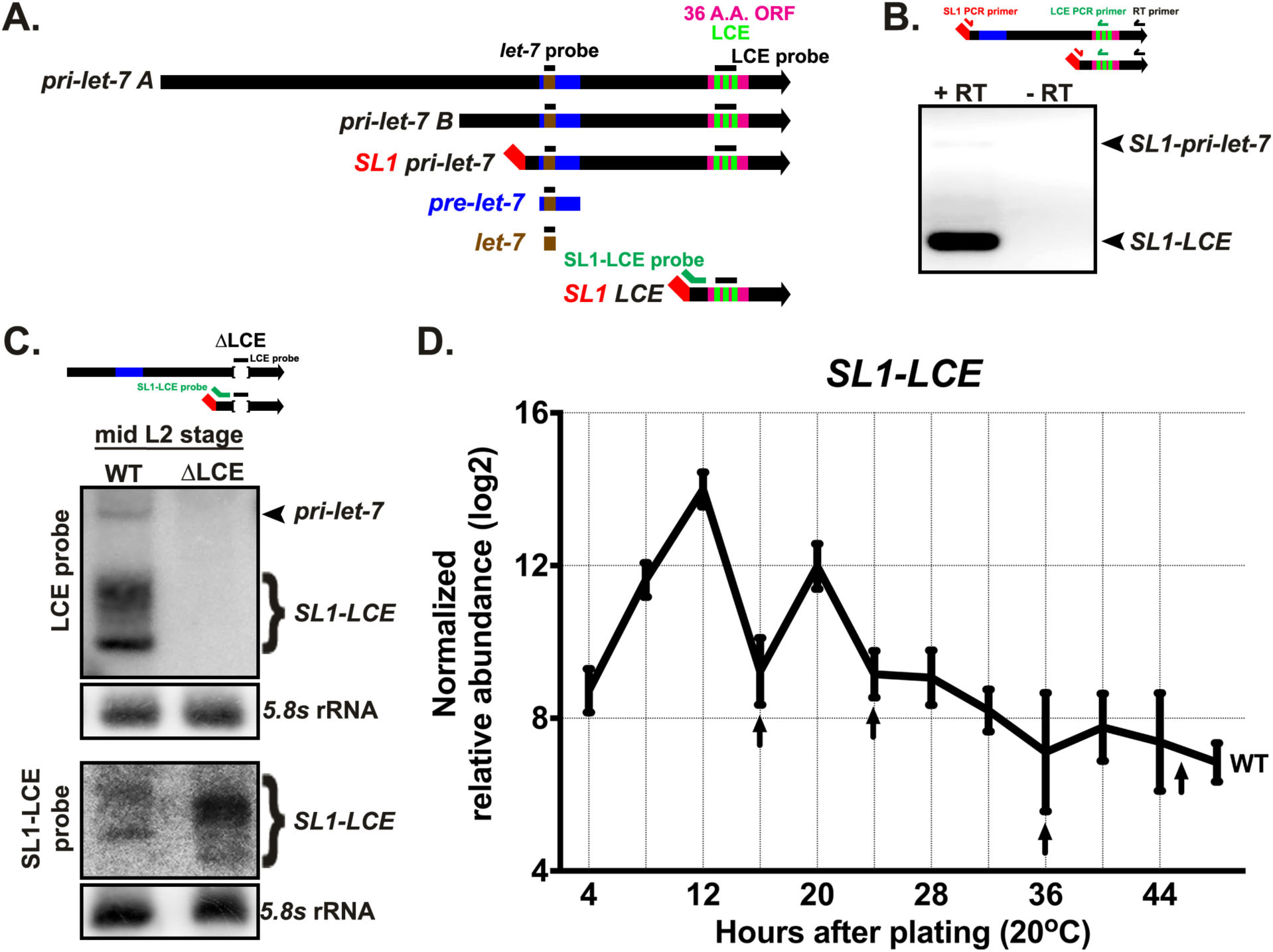
The *C. elegans let-7* locus produces four transcripts. (A) Major transcripts produced from the *C. elegans let-7* locus, and the locations of probes used in Northern blots shown in D, and in subsequent Figures. (B) Non-qRT-PCR analysis of total RNA from a mix-population of wild-type (WT) animals, using an SL1 forward primer and LCE reverse primer, with (+) and without (-) reverse transcriptase (RT) in the cDNA synthesis reaction. (C) Total RNA from WT and LCE deletion mid-L2 animals (20 hours after plating) analyzed by northern blotting with probes for the LCE and SL1-LCE. (D) qRT-PCR analysis of SL1-LCE levels in samples of total RNA from staged populations of synchronously developing WT animals. Data are represented as mean ±SD. Arrows mark the times of larval molts.

*pri-let-7* transcripts are expressed at all four larval stages of *C. elegans* development (L1-L4), whereas mature *let-7* is abundantly expressed only in the L3 and L4 larval stages. Intriguingly, *pri-let-7* levels oscillate during each larval stage, peaking mid-stage and dipping during larval molts, likely as a result of underlying pulsatile transcriptional activity of the *let-7* locus (Van Wynsberghe et al., 2011, McCulloch and Rougvie, 2014, Perales et al., 2014, Van Wynsberghe et al., 2014). Why *pri-let-7* pulses with each larval stage remains unclear. However, the dramatic discrepancy between *pri-let-7* and mature *let-7* levels at early larval stages indicates potent post-transcriptional inhibition of *let-7* biogenesis during the L1 and L2 stages; indeed, the RNA-binding protein, LIN-28, is expressed at early larval stages in *C. elegans* and exerts a strong inhibition of *let-7* processing (Van Wynsberghe et al., 2011). While CLIPseq data indicated that LIN-28 could bind *pri-let-7* in *vivo* to inhibit accumulation of mature *let-7* microRNA (Stefani et al., 2015), the precise mechanisms by which LIN-28 exerts this inhibition of *let-7* biogenesis remain unclear.

Trans-splicing is the act of joining two separate RNAs. In *C. elegans*, approximately 70% of all transcripts (including mRNAs and microRNA primary transcripts) are trans-spliced with a 22-nucleotide splice leader RNA. The outcomes of trans-splicing include separating the individual mRNAs of polycistronic operon transcripts, shortening the 5’ untranslated regions (5’ UTRs) of RNAs, and changing the 5’ RNA cap from monomethyl to trimethyl guanosine. There are two SL RNAs with distinct sequences: SL1 and SL2 (of which there are 11 variants). While there are exceptions, SL1 trans-splicing tends to be the 5’ most trans-splicing event of a primary RNA transcript and invariably results in rapid degradation of the 5’ “outron”. SL2 trans-splicing is generally restricted to downstream ORFs of operons after SL1 trans-splicing and polyadenylation of the first mRNA (Morton and Blumenthal, 2011, Blumenthal, 2012). Similar to cis-splicing, trans-splicing in *C. elegans* uses a consensus acceptor sequence of “TTTCAG” (Graber et al., 2007).

Here we identify a previously-undescribed trans-splicing event in *pri-let-7* that occurs downstream of the *let-7* stem-loop, and produces a short (approx. 262nt) mRNA that contains a 5’ SL1 leader sequence, a short ORF, *let-7* complimentary sequences, and a poly-A tail. We provide evidence that LIN-28 is necessary for this splicing event, and that trans-splicing serves to negatively regulate *let-7fam* in two ways: First, by preventing precocious *let-7* expression through the degradation of the upstream outron which contains the *let-7* precursor; second, by inhibiting *let-7fam* activity via production of an RNA that functions as a *let-7fam* sponge. Thus, we have characterized a splicing event involving *let-7* primary transcripts that can regulate *let-7* biogenesis in *cis* as well as *let-7fam* activity in *trans*.

## RESULTS

### The *let-7* locus produces a short trans-spliced transcript that contains *let-7* complimentary sequences

As mentioned above, a previous study identified a region of *pri-let-7* in *C. elegans* that contains a *let-7* <u>c</u>omplimentary <u>e</u>lement (LCE) (Zisoulis et al., 2012). We confirmed that three sites within the LCE element have complementarity to *let-7fam* microRNAs so as to permit base pairing to the *let-7*-family seed sequence plus varying degrees of 3’ supplemental pairing (Fig. S1a). We also noted that an additional transcript, C05G5.7, is annotated to be transcribed from the *let-7* locus and contains a 647 nt 5’ UTR that contains the *let-7* stem-loop, a 111 nt ORF that contains the LCE, and a 79 bp 3’ UTR. We also identified a potential splice acceptor sequence located upstream of the LCE and downstream of the *let-7* stem-loop, which could mediate SL1-trans-splicing and thereby produce a short ∼262nt transcript containing the LCE but lacking the upstream *let-7* stem-loop (Fig. 1A, Fig. S1A). Analysis of the *let-7* locus of other *Caenorhabditis* species showed both the trans-splice acceptor sequence and the LCE are conserved (Figure S1A). To determine if an ∼262 nt SL1-spliced LCE transcript is expressed *in vivo*, we performed non-quantitative PCR (non-qRT-PCR) using SL1 forward and LCE reverse primers, and we observed two distinct bands. We determined the top band to be the previously known SL1-spliced version of *pri-let-7* (with the SL1 upstream of the *let-7* stem-loop), and the bottom band to be the predicted SL1-spliced LCE transcript (hereafter referred to as *SL1-LCE*) (Fig. 1B). We also determined that both *SL1-LCE* and *SL1-pri-let-7* are poly-adenylated, by generating cDNA using oligo-dT, followed by non-qRT-PCR (Fig. S2).

We confirmed the existence of *SL1-LCE* by northern blotting with a probe to the LCE sequence and determined that *SL1-LCE* is more abundant than *pri-let-7* at the mid-L2 stage (Fig. 1C). Interestingly, we observed a range of *SL1-LCE* lengths, from an estimated 275 to 340 nt. After probing specifically to the SL1 splice junction at the 5’ end of *SL1-LCE*, we observed the same range in length, indicating that this variation is at the 3’ end, and suggests either differences in poly-adenylation or transcriptional stop sites (Fig. 1C). To confirm that the variable length was not at the 5’ end, we also performed 5’RACE and observed only a single 5’ terminus, at the site of the *SL1-LCE* trans splice (Fig. S3). We also confirmed the three 5’ ends of *pri-let-7* (Fig. S3 and data not shown). We failed to detect the 5’ end of the annotated transcript C05G5.7

The location of the trans-splice acceptor sequence within the *let-7* locus suggests that *SL1-LCE* could be processed from *pri-let-7*. Therefore, we tested whether the expression pattern of *SL1-LCE* is also developmentally regulated, like *pri-let-7*. Using quantitative RT-PCR (qRT-PCR), we determined that similar to *pri-let-7, SL1-LCE* levels pulsed in phase with the cycle of larval molts, indicating that *SL1-LCE* expression could be driven by the same oscillatory transcriptional program as *pri-let-7* (Van Wynsberghe et al., 2011, Perales et al., 2014). However, unlike *pri-let-7*, *SL1-LCE* stopped pulsing and remained relatively low after the L2 stage (Fig. 1D). These finding suggest that *SL1-LCE* is generated from *pri-let-7* by trans-splicing, but that it is post-transcriptionally down regulated in conjunction with larval developmental progression.

### *lin-28* regulates the expression of *SL1-LCE*

Coordinated with a variety of other factors, *let-7* functions within the heterochronic pathway to ensure proper developmental timing of various cell fates, in particular progression from larval to adult fates during the L4-to-adult transition (Reinhart et al., 2000). The high L1 and L2 expression of *SL1-LCE* suggested that the heterochronic genes that specify early larval events could promote the expression of *SL1-LCE.* Therefore, we assessed *SL1-LCE* expression in heterochronic mutants with altered temporal patterns of early larval cell fates.

*lin-4* is a microRNA that is necessary for the transition from the L1 to L2 stages through its targeted repression of the *lin-14* 3’UTR. *lin-4* null (*lin-4(0)*) mutations, or *lin-14* gain of function (*lin-14(gf)*) mutations, result in aberrant up regulation of *lin-14* that retards developmental progression by continuously specifying the repetition of L1 events (Chalfie et al., 1981, Ambros and Horvitz, 1984, Ambros, 1989, Lee et al., 1993). On the other hand, *lin-14* loss-of-function (*lin-14(lf)*) results in premature developmental progression, characterized by a skipping of L1 events and precocious advancement through the L2 to adult stages (Ambros and Horvitz, 1984). We observed that *SL1-LCE* levels remained high throughout development in *lin-4(0)* animals and also in *lin-14(gf)* animals, while *SL1-LCE* levels were unchanged in *lin-14(lf)* (Fig. 2A) indicating that *lin-14* promotes, but is not necessary for *SL1-LCE* expression.

**Figure 2.**
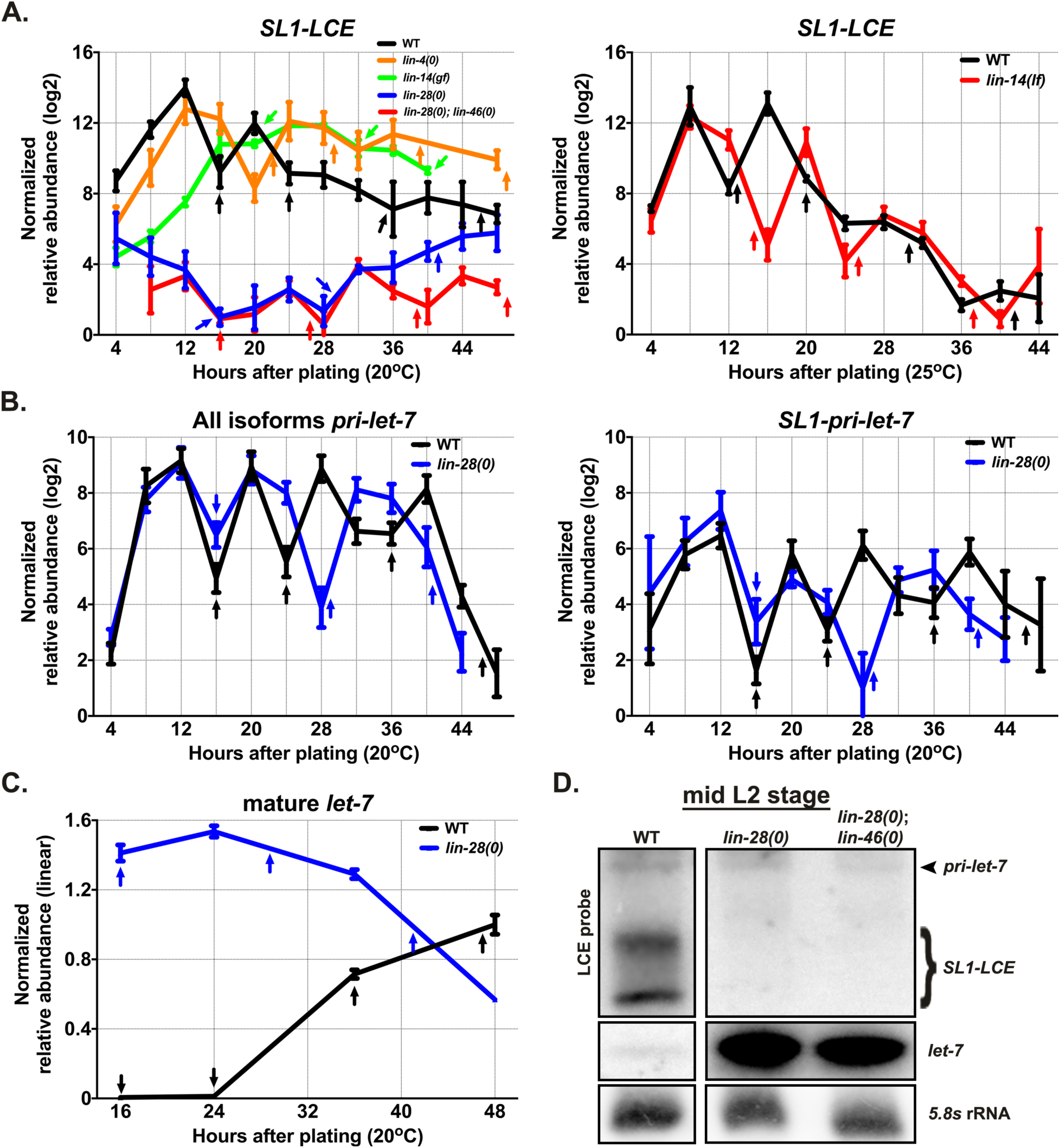
*lin-28* is necessary for the expression of *SL1-LCE*. (A) qRT-PCR analysis of SL1-LCE levels in samples of total RNA from staged populations of synchronously developing WT animals, and various heterochronic mutants. Data are represented as mean ±SD. Arrows mark the times of larval molts. (B) qRT-PCR analysis of the levels of all *pri-let-7* isoforms (left panel) and *SL1-pri-let-7* (right panel) in samples of total RNA from staged populations of synchronously developing WT animals, and *lin-28(0)* animals. Error bars represent SDs. Arrows mark the times of larval molts. (C) FirePlex miRNA analysis of *let-7* levels in samples of total RNA from staged populations of synchronously developing WT animals and *lin-28(0)* animals. Data are represented as mean ±SD. Arrows mark the times of larval molts. (D) Total RNA from WT, *lin-28(0)*, and *lin-28(0)*; *lin-46(0)* mid-L2 animals (20 hours after plating) analyzed by northern blotting, and hybridized with probes for either the LCE (top two panels) or for *let-7* (middle two panels).

*lin-28* is an RNA binding protein that functions in early larval stages to regulate developmental progression from L2 to later cell fates. Loss of *lin-28* results in the skipping of L2 events, precocious advancement to the adult stage, and precocious expression of *let-7* (Fig. 2A and 2D), (Ambros and Horvitz, 1984, Van Wynsberghe et al., 2011). We found that *lin-28(0)* animals exhibited drastically reduced *SL1-LCE* levels compared to the wild-type (Fig. 2A and 2D). Therefore, in contrast to *lin-14, lin-28* seems to be essential for expression of the *SL1-LCE* transcript.

The fact that *lin-14(lf)* animals have precocious phenotypes like those of *lin-28(0*), yet have normal *SL1-LCE* expression, argues that the reduced *SL1-LCE* expression in *lin-28(0)* is not an indirect consequence of precocious development. In further support of this conclusion, we observed that in *lin-28(0);lin-46(0)* animals, which are completely suppressed for precocious phenotypes but not for precocious *let-7* levels (Pepper et al., 2004, Vadla et al., 2012), (Fig. 2D), *SL1-LCE* levels are similarly reduced as in *lin-28(0)* alone (Fig. 2A and 2D). Taken together, this suggests that *lin-28* has a relatively direct role in promoting LCE trans-splicing.

One possible explanation of why *SL1-LCE* levels are low in *lin-28(0)* is that expression of *pri-let-7* could be reduced. To test this possibility, we measured the levels of *pri-let-7* using qRT-PCR and observed no reduced expression of *pri-let-7* in *lin-28(0)* larvae (Fig. 2B and D). This finding argues that LIN-28 post-transcriptionally regulates the generation of *SL1-LCE*. We also noted that despite the fact that *lin-28(0)* animals undergo only three larval stages instead of the normal four, the pulses of *pri-let-7* still coincided with each larval molt (Fig. 2B).

As mentioned previously, one *pri-let-7* isoform is SL1 trans-spliced upstream of the *let-7* stem-loop. To test if loss of *lin-28* loss of function also reduced this upstream trans-splicing event, we measured *SL1-pri-let-7* levels in *lin-28(0)* animals and observed no significant reduction (Fig. 2B). Put together, these data indicate that LIN-28 is essential for generating *SL1-LCE* from *pri-let-7* by specifically promoting trans-splicing at the downstream splice acceptor.

### Mutations of the LCE splice acceptor result in the use of cryptic splice acceptor sequences

To determine the function of trans-splicing of the LCE, we used CRISPR/Cas9 to introduce mutations of the “TTTCAG” SL1 acceptor sequence of the LCE transcript. To our surprise, deletion of the TTTCAG sequence did not eliminate LCE trans splicing; non-qRT-PCR and TA-cloning revealed that SL1 trans-splicing still occurred using a cryptic acceptor sequence (TTGTAG) located 27 nucleotides upstream of the canonical TTTCAG (data not shown). qRT-PCR revealed that in wild-type animals use of this cryptic TTGTAG sequence was minimal (Fig. S4A), whereas in the animals in which the canonical TTTCAG was deleted, use of the cryptic TTGTAG was readily detectable and the expression pattern of the resulting (albeit slightly longer) *SL1-LCE* was similar to that of the normal *SL1-LCE* in wild-type animals (Fig. S4A). Furthermore, the use of this cryptic TTGTAG splice acceptor was dependent upon *lin-28* (Fig. S4A) indicating that LIN-28 can promote LCE-proximal trans-splicing regardless of the acceptor sequence.

When we deleted the canonical TTTCAG, northern blots showed a decrease in *SL1-LCE* levels and an increase in *pri-let-7* levels (Fig. 3B) signaling that *SL1-LCE* is trans-spliced from *pri-let-7*, and the non-canonical TTGTAG SL1 acceptor sequence is not as efficient in this context as is the wild-type TTTCAG sequence. Moreover, because this cryptic TTGTAG sequence is upstream to the canonical TTTCAG, all of the corresponding *SL1-LCE* bands were shifted up strengthening our interpretation that all northern blot bands associated with this transcript are SL1 trans-spliced (Fig. 3B).

**Figure 3.**
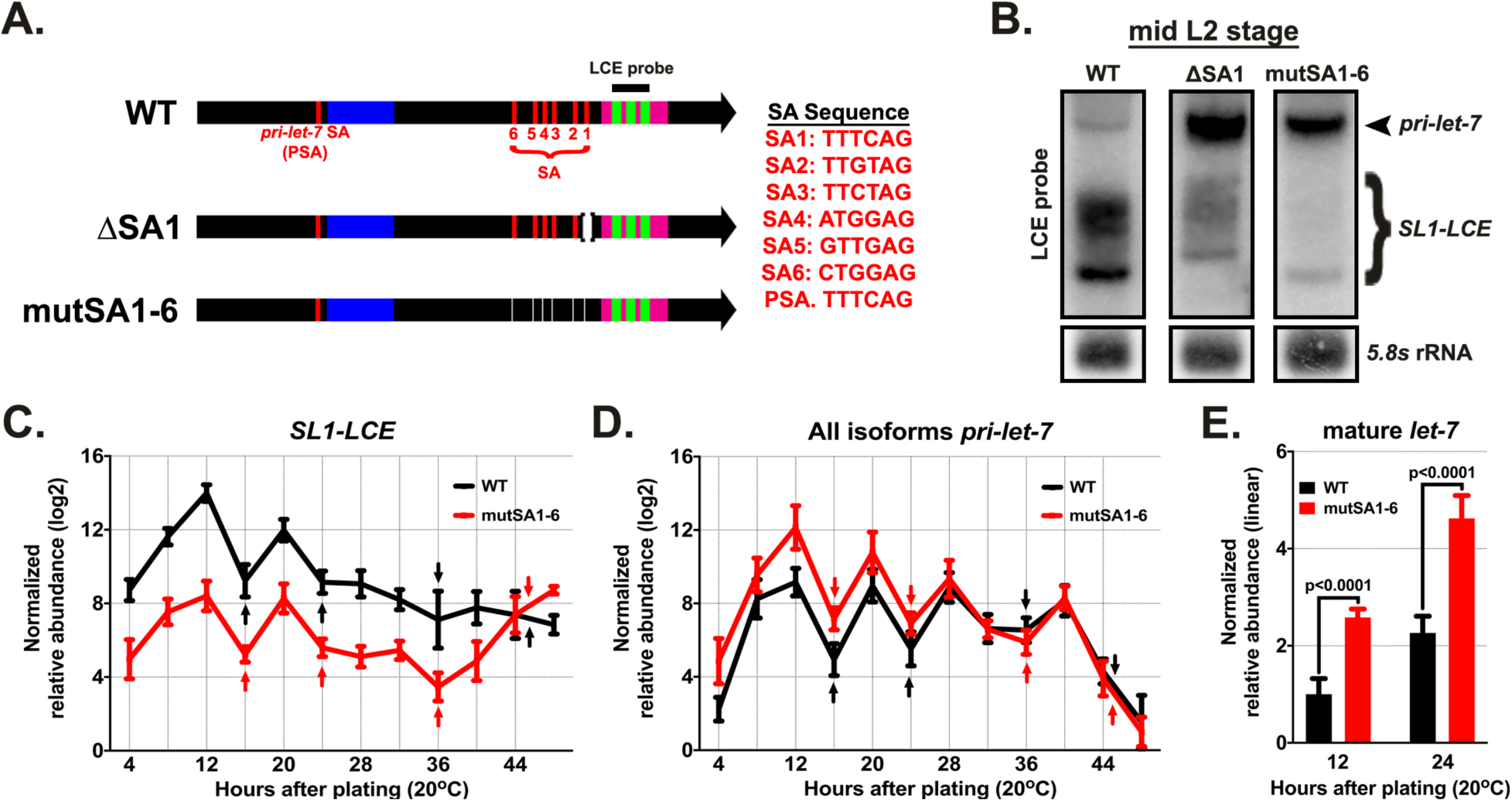
Larvae with *SL1-LCE* splice acceptor site mutations display elevated levels of *pri-let-7* and mature *let-7.* (A) Locations of the NTNNAG motif SL1 splice acceptors (SA) in the *let-7* locus and the SA mutations used in these experiments. PSA refers to the splice acceptor sequence in *pri-let-7* upstream of the *pre-let-7* stem-loop. SA1 is the position of trans splicing that generates *SL1-LCE* in wild-type. SA2-SA6 are non-canonical splice acceptor sequences, some which can be utilized when SA1 is mutated. (B) Total RNA from mid-L2 (20 hours after plating) WT, SA1 deletion, and mutSA1-6 animals, analyzed by northern blotting, hybridized with a probe for the LCE. (C) qRT-PCR analysis of the levels of *SL1-LCE* in samples of total RNA from staged populations of synchronously developing WT animals, and mutSA1-6 animals. Data are represented as mean ±SD. Arrows mark the times of larval molts. (D) qRT-PCR analysis of the levels of all *pri-let-7* isoforms in samples of total RNA from staged populations of synchronously developing WT animals, and mutSA1-6 animals. Data are represented as mean ±SD. Arrows mark the times of larval molts. (E) FirePlex miRNA analysis of *let-7* levels in in samples of total RNA from staged populations of synchronously developing WT and mutSA1-6 animals at the mid-L1 (12 hours after plating) and mid-L2 (20 hours after plating) stages. Error bars represent SDs. Statistical significance was determined using a Student’s t test.

Based on previous genomic analysis of SL1 splice sites, the consensus SL1 acceptor sequence is TTTCAG. However, other acceptor sequences can be utilized but certain nucleotides appear to remain invariant, namely a T in the second position, A in the fifth position, and G in the sixth position (Graber et al., 2007). There are six occurrences of the corresponding NTNNAG consensus sequence in the region between the *let-7* stem-loop and the LCE (Fig. 3A). With the aim of eliminating trans splicing altogether in this region, we made small deletions in all six SAs using CRISPR/Cas9. Non-qRT-PCR and sequence analysis of LCE transcripts from the six-fold SA mutant (mutSA1-6) revealed that some basal level of trans-splicing still occurred, now using two far-non-canonical acceptor sequences, TTTCGG and TTCGGG, which are one nucleotide apart from each other and near the original splice site location (data not shown). Compared to wild-type, qRT-PCR and northern blotting of mutSA1-6 showed a drastic reduction in *SL1-LCE* levels as well as an increase in *pri-let-7* in the L1 and L2 stages (Fig. 3C and 3D).

### *SL1-LCE* trans-splicing regulates *let-7* processing

In *C. elegans,* the 5’ region of an RNA that is removed by SL1 trans-splicing is called the outron. The outrons of trans-spliced RNAs are rarely detected, suggesting they are rapidly degraded following trans-splicing (Morton and Blumenthal, 2011). Previously published northern blots of *pri-let-7* suggest that the outron is below detectable levels (Bracht et al., 2004, Van Wynsberghe et al., 2011, Zisoulis et al., 2012, Van Wynsberghe et al., 2014). The resolution of our northern blots could not definitively show if the outron was detectable, so we employed qRT-PCR using RT primers positioned 5’ or 3’ of the *SL1-LCE*’s splice acceptor, and PCR primer pairs flanking the *SL1-LCE* splice acceptor. cDNA primed from sequences 5’ of the splice acceptor would represent both the pre-spliced *pri-let-7* as well as the post-spliced outron, whereas cDNA primed from sequences 3’ of the splice acceptor would represent only pre-spliced *pri-let-7*. Therefore, if the outron were present at detectable levels, cDNA from sequences 5’ of splice acceptor would more abundant than cDNA from sequences 3’ of the splice acceptor.

When we performed qRT-PCR to *pri-let-7*, we observed no difference in the yields of cDNA from sequences 5’ versus 3’ of the splice acceptor. This indicates that the outron is undetectable by this assay, and hence relatively unstable compared to unspliced *pri-let-7* (Fig. S4B).

If trans splicing of *SL1-LCE* produces an unstable outron containing unprocessed *let-7* microRNA, we hypothesized that trans-splicing of *SL1-LCE* from *pri-let-7* could have a net negative effect on accumulation of mature *let-7* microRNA. In L1 and L2 animals when trans-splicing is normally abundant, *let-7* microRNA levels in mutSA1-6 animals were approximately twice as high compared to wild-type, indicating that trans-splicing of *SL1-LCE* negatively impacts accumulation of mature *let-7* (Fig. 3E and S4C). Interestingly, at later developmental stages when *SL1-LCE* trans-splicing is not prevalent, *let-7* levels were reduced by ∼60% to 70% suggesting an additional positive regulatory role for sequences overlapping one or more of the SA elements mutated in mutSA1-6 (Fig. S4D).

### The LCE functions to negatively regulate the *let-7* family

A previous study reported experiments suggesting that the LCE region in *pri-let-7*, in conjunction with the Argonaut protein ALG-1, could function in *cis* to facilitate *let-7* biogenesis (Zisoulis et al., 2012). In that study, a transgene carrying a modified *let-7* locus with a deletion of 178bp (which removed the LCE and surrounding sequences) expressed decreased levels of mature *let-7*, suggesting that the LCE, perhaps when bound to *let-7* miRISC, could function to promote microprocessing of *pri-let-7*. However, because this 178bp deletion also removed sequences upstream of the LCE, including the SL1-acceptor sequence, we used CRISPR/Cas9 to create a 55bp deletion at the endogenous *let-7* locus that removed only the LCE. This 55bp deletion of the LCE did not result in a measurable change in *let-7* levels (Fig 4B). We also used CRISPR/Cas9 to introduce the previously described 178bp deletion at the endogenous *let-7* locus, and confirmed the previous (Zisoulis et al., 2012) results: an approximately ten-fold reduction *let-7* levels (Fig. 4B). Based on these results, we conclude that the larger 178bp deletion removes unknown non-LCE positive regulatory elements, and that the LCE sequences themselves do not exert a detectable positive effect on *let-7* biogenesis.

**Figure 4.**
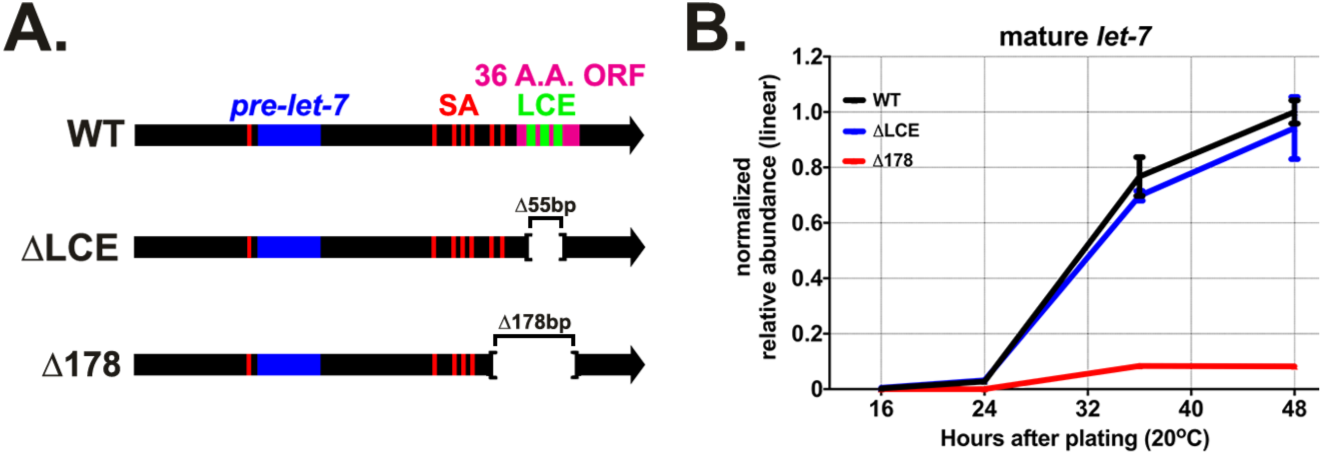
LCE sequences are dispensable for normal expression of mature *let-7* microRNA. (A) Positions of the *let-7* mutants used in these experiments. (B) FirePlex miRNA analysis of *let-7* levels in WT, LCE deletion (ΔLCE), and animals containing a 178bp deletion (Δ178) previously reported to reduce *let-7* biogenesis throughout development. Data are represented as mean ±SD.

Because *SL1-LCE* contains sequences that are complimentary to *let-7fam*, we hypothesized that *SL1-LCE* could function as a sponge to negatively regulate *let-7fam*. To test this, we sought to determine if mutations disrupting *SL1-LCE* could genetically interact with *let-7fam* in sensitized genetic backgrounds. Of the four major *let-7* family genes, only *let-7(lf)* or *mir-48(0)* mutants display overt heterochronic phenotypes in normal laboratory conditions. Loss of either *mir-48* or *let-7* results in retarded hypodermal development and an extra larval molt. Additionally, *let-7(lf)* hermaphrodites burst through an improperly formed vulva (an adult lethality phenotype), and *mir-48(0)* adult hermaphrodites die due to egg retention as a result of their retarded hypodermal development. We hypothesized that if *SL1-LCE* were to function as a sponge for the *let-7fam* microRNAs, loss of the LCE sequences would result in increased *let-7fam* activity, which could be evidenced by suppression of *let-7(lf)* or *mir-48(0)* phenotypes.

Col-19::GFP is a reporter that is expressed in hypodermal seam cells and in the hypodermal syncytium (hyp-7) beginning at the L4 molt. In *let-7(lf)* as well as *mir-48(0)* animals at the L4 molt, Col-19::GFP expression is dramatically reduced in hyp-7 compared to the wild-type, and is limited to the seam cells. Using CRISPR/Cas9 we introduced a deletion of the *let-7* hairpin into the LCE deletion background and observed no suppression of the *let-7(lf)* phenotypes: vulva bursting and reduced Col-19::GFP expression were unchanged compared to *let-7(lf)* with the LCE intact (data not shown). Therefore, absence of the LCE did not display any evident genetic interaction with *let-7(lf)*. By contrast, a strong genetic interaction was evident between LCE deletion and *mir-48(0)*.

When the LCE deletion mutation was crossed into *mir-48(0)*, col-19::GFP expression was restored to a wild-type pattern (Fig. 5). This suggests that removal of the LCE sequences results in up regulation of the activity of one or more members of *let-7fam*. Consistent with this supposition, when a second family member, *mir-241*, was also removed, loss of the LCE failed to restore the normal timing of col-19::GFP expression (Fig. 5B). Deletion of the LCE similarly suppressed the extra molting phenotype, restored survival, and restored the brood size of *mir-48(0)* while only partially doing so in *mir-48(0), mir-241(0)* double nulls (Fig. S5A-C). To determine if this suppression was due to an elevation in *let-7fam* levels, we measured their respective levels in wild-type and LCE deletion animals and observed no difference (Fig. S6A-C). Put together these results indicated that the LCE negatively regulates *let-7fam* by modulating their activity rather than their levels.

**Figure 5.**
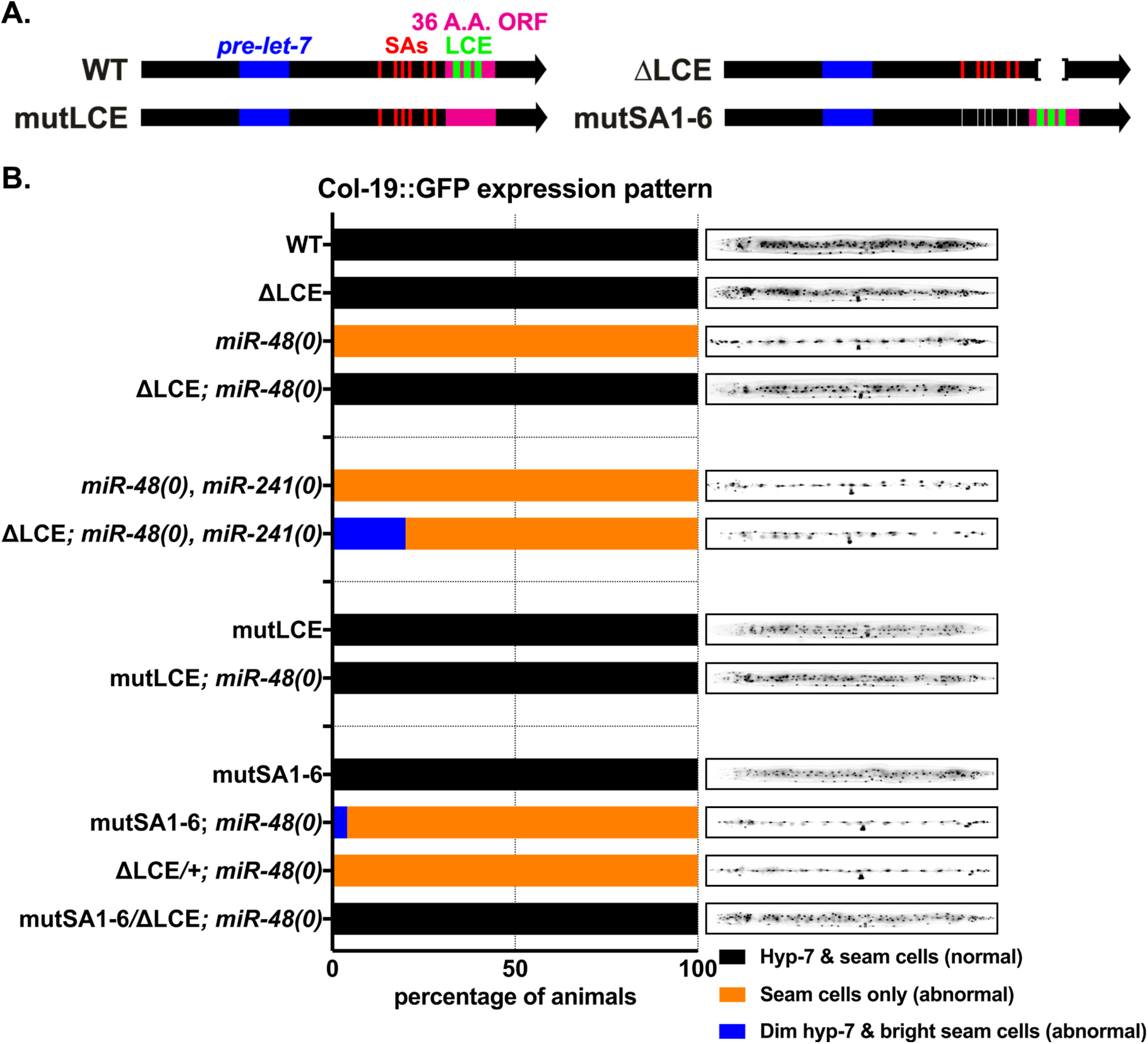
Inhibition of *SL1-LCE* function, either by deletion of the LCE, or by mutations of LCE-proximal trans-splicing acceptor sequences, suppresses the retarded hypodermal phenotypes of *mir-48(0)* animals. (A) Positions of the *let-7* LCE mutations (green) and SA mutants (red) used in these experiments. (B) Deletion or mutation of the LCE or mutation in the SAs suppresses *mir-48(0).* Col-19::GFP phenotype in molting late L4 animals. The graph is a quantification of the Col-19::GFP phenotypes observed for each genotype. N ≥ 9. Images are of a representative molting L4 animal for each genotype.

The LCE sequence is predicted to contain a 36 amino acid ORF that is poorly conserved in other *Caenorhabditis* species (Fig. S1A), and it is therefore unlikely to perform a conserved function. Nevertheless, our 55bp deletion of the LCE disrupts this putative ORF so it was possible that disruption of the ORF could confound the interpretation of our results. Therefore we used CRISPR/Cas9 to mutate the three *let-7* complimentary sequences (LCSs) without altering the amino acid sequence of the ORF. Similar to the LCE deletion, these ‘silent’ LCS mutations did not alter *let-7* levels (data not shown). However, these LCE mutations did restore normal Col-19::GFP expression, normal survival, normal brood size, and suppressed the extra molting phenotype in *mir-48(0)* animals but not in *mir-48(0), mir-241(0)* double null animals (Fig. 5B and Fig. S5A-C).

### Tran-splicing of the LCE is necessary to negatively regulate *let-7 family* microRNAs

To function as a negative regulator of *let-7fam* activity by acting as a microRNA sponge, the *SL1-LCE* transcript would presumably encounter *let-7fam* miRISC in the cytoplasm. As mentioned above, *SL1-LCE* contains a putative 36 amino acid ORF. Previously published ribosome-profiling data indicated that ribosomes locate to the LCE sequence (Michel et al., 2014), suggesting the LCE ORF is translated, and therefore enters the cytoplasm where it could engage *let-7 family* microRNAs. To confirm that the LCE ORF can be translated *in vivo*, we generated transgenic animals carrying a transgene with the C-terminus of the LCE ORF fused to GFP, and observed fluorescence that recapitulated the temporal expression of *SL1-LCE* (Fig. S7).

In addition to being cytoplasmic, to function as a sponge, the *SL1-LCE* transcript should presumably be expressed at levels in molar excess to *let-7fam* microRNA*s*. To determine the stoichiometric ratio of *SL1-LCE* to *let-7fam* in wild-type larvae, we quantified the amount of *SL1-LCE* and *let-7fam* microRNAs in RNA samples from synchronized populations of developing larvae. We calibrated these assays using known amounts of *in vitro* transcribed *SL1-LCE* and *pri-let-7,* and synthetic *let-7fam* microRNA oligonucleotides. The results of these quantitative assays indicated that the *SL1-LCE* is in molar excess to *let-7*, *mir-48, mir-84,* and *mir-241* during the L1 and L2 stages (Fig. 6A). Importantly, *pri-let-7* levels were not in excess to *let-7fam,* indicating that *pri-let-7*, despite containing LCE sequences, is not likely to contribute as significantly to *let-7fam* sponging as does the *SL1-LCE* transcript.

**Figure 6.**
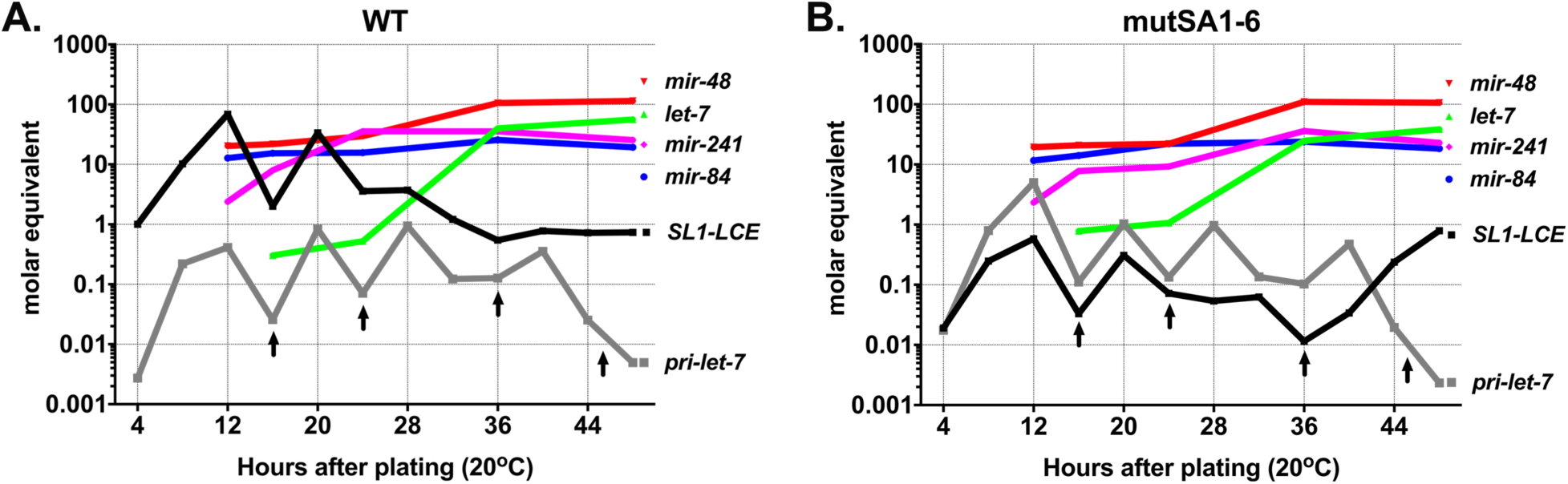
The *SL1-LCE* is in molar excess relative to *let-7fam* microRNAs during early larval stages. Relative levels (molar equivalents, normalized to *in vitro* transcribed *pri-let-7* and synthetic *let-7fam*) of *pri-let-7*, *SL1-LCE*, *mir-48*, *mir-84*, *mir-241*, and *let-7,* determined by calibrated qRT-PCR (for *pri-let-7* and *SL1-LCE*), or calibrated Fireplex assay (for microRNAs), in samples of total RNA from staged populations of synchronously developing WT animals (A), and mutSA1-6 animals (B). Arrows mark the times of larval molts.

**Figure 7.**
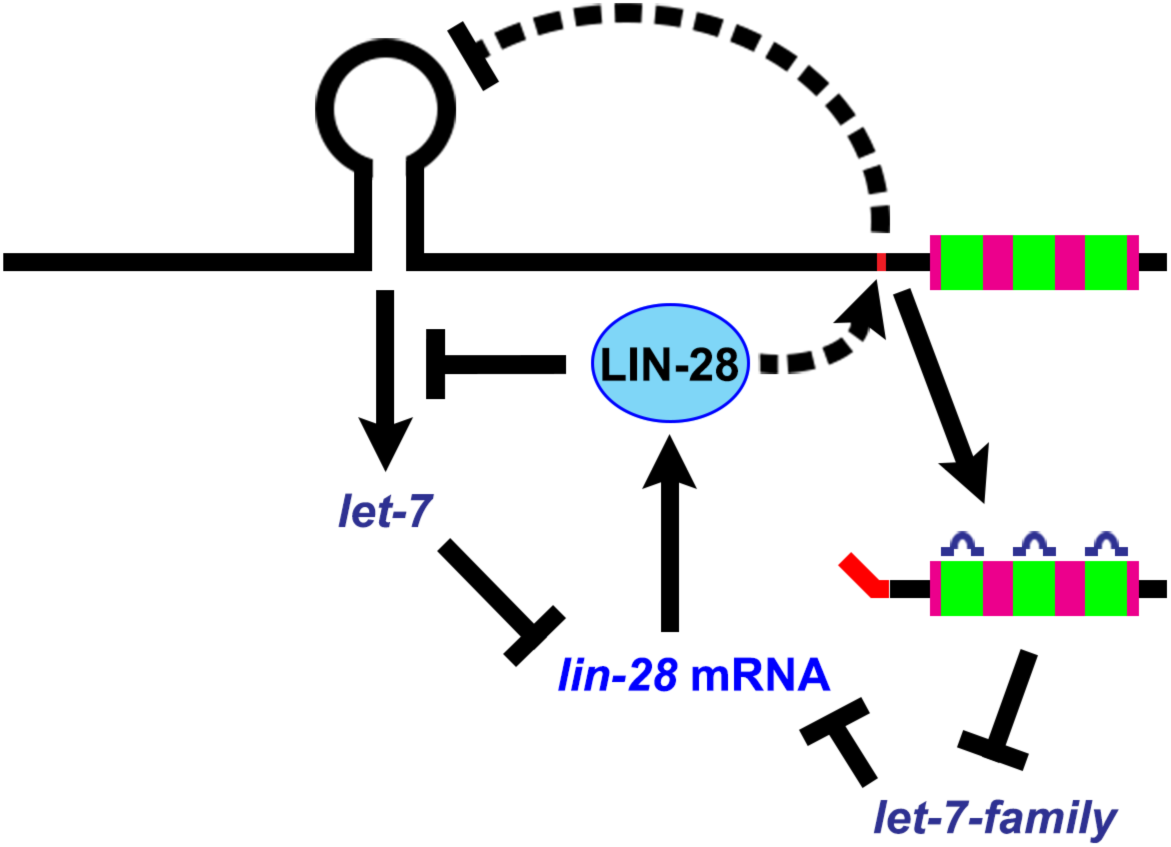
A model of how LIN-28 and *let-7fam* microRNAs function in a reciprocal negative regulatory network. A summary of the genetic pathway determined by our results in which *lin-28* and *let-7fam* post-transcriptionally exert reciprocal negative feedback on each other.

Based on its relative abundance and cytoplasmic location, our results suggest that of the two classes of LCE-containing transcripts produced from the *let-7* locus (*pri-let-7* and *SL1-LCE*), the *SL1-LCE* has the ability to function as a sponge for *let-7fam* microRNA. This thereby advocates that the sponging activity of the LCE would depend on trans splicing of *SL1-LCE*. To test this supposition, we took advantage of the splice site (mutSA1-6) mutant animals, which have reduced *SL1-LCE* levels. Using the same calibrated quantitation as above, we determined that the reduced *SL1-LCE* levels in mutSA1-6 animals were not in molar excess of *let-7fam* (Fig. 6B). Moreover, the elevation in *pri-let-7* in the mutSA1-6 was not sufficient to put it in molar excess of all *let-7fam* (Fig. 6B), although we note that in mutSA1-6, *pri-let-7* was in excess to *mir-241* and *let-7* at the L1 peak and approximately equimolar with *let-7* at the L2 peak (Fig. 6B).

We hypothesized that, similar to deletion of the LCE sequences, the reduction in *SL1-LCE* levels in mutSA1-6 would increase the activity of *let-7fam* and suppress *mir-48(0)* phenotypes. However, when mutSA1-6 and *mir-48(0)* were combined, we observed no suppression (Fig. 5B and S5A-C). This suggested that although mutSA1-6 causes a significant decrease in *SL1-LCE,* the remaining LCE containing transcripts could be nevertheless functional. Therefore, we aimed to reduce the amount of remaining *SL1-LCE* in half by using mutSA1-6/ΔLCE trans-heterozygous animals. This further reduction in *SL1-LCE* resulted in suppression of all the heterochronic phenotypes associated with *mir-48(0),* supporting the conclusion that *SL1-LCE* functions to negatively regulate *let-7* family activity (Fig. 5B and S5A-C).

## DISCUSSION

A general property of microRNAs across diverse organisms is that they are first transcribed as longer primary transcripts, which are then enzymatically processed to produce the mature 22nt microRNA. The multiple biogenesis steps required to generate a mature microRNA provide access for a range of transcriptional and post-transcriptional regulation. For example in *C. elegans*, HBL-1 and LIN-42 can modulate the transcriptional activity of microRNAs including *let-7* and *lin-4* (Roush and Slack, 2009, McCulloch and Rougvie, 2014, Perales et al., 2014, Van Wynsberghe et al., 2014), and LIN-28 post-transcriptionally regulates *let-7* (Van Wynsberghe et al., 2011). The negative regulation of *let-7* by LIN-28 is conserved evolutionarily. In mammals, it has been shown that LIN28 can bind to the stem-loop of *pri-let-7* and/or *pre-let-7* to directly inhibit processing by Drosha and/or Dicer (Heo et al., 2008, Newman et al., 2008, Rybak et al., 2008, Viswanathan et al., 2008, Heo et al., 2009, Loughlin et al., 2011, Nam et al., 2011, Piskounova et al., 2011). In *C. elegans,* LIN-28 also appears to inhibit processing of *pri-let-7* by Drosha, although apparently not by binding the *let-7* stem loop, but rather through binding to sequences ∼170 nucleotides downstream (Stefani et al., 2015). Also in both *C. elegans* and mammals, the 3’ UTR of *lin-28* contains sequences complimentary to the *let-7fam* indicating that *let-7fam* can repress LIN-28 expression (Reinhart et al., 2000, Rybak et al., 2008). Thus, *lin-28* engages in an evolutionarily conserved reciprocal negative feedback with *let-7family* microRNAs. However, the functional significance of these *lin-28-let-7fam* regulatory interactions, and their precise mechanisms, are not fully understood.

In this study, we identified a previously undescribed RNA, *SL1-LCE*, which is trans-spliced from *C. elegans pri-let-7* downstream of the *pre-let-7* stem-loop, and that contains *let-7* complimentary sequences followed by 3’ poly-A. We determined that *SL1-LCE* is highly expressed in the early larval stages, displaying an inverse expression pattern compared to *let-7,* suggesting that it could be associated with negative regulation of *let-7* biogenesis. We find that expression of *SL1-LCE* coincides with the expression of LIN-28, and is dependent on *lin-28* function, revealing a novel regulatory circuit in which LIN-28 governs trans-splicing of *pri-let-7* to negatively impact *let-7* microprocessing. The regulation of *SL1-LCE* trans-splicing by *lin-28* appears to be independent of other phenotypes controlled by *lin-28*, as another precocious mutant *lin-14(lf)* had no affect on trans-splicing, and *SL1-LCE* levels were low in *lin-28(0);lin-46(lf)*, where precocious phenotypes are suppressed.

Interestingly, when we mutated the *SL1-LCE* trans-splice acceptor sequence, even in combination with mutations of nearby putative splice acceptors, trans-splicing persisted using a far-non-canonical acceptor sequences. This suggests the presence sequences in *pri-let-*7 with potent splicing enhancer activity. While the use of a far-non-canonical sequence is unusual, it is not unprecedented in *C. elegans* (Aroian et al., 1993).

Our findings suggest that LIN-28 inhibits *let-7* biogenesis through the combined effects of two mechanisms. On the one hand, LIN-28 binds directly to *pri-let-7* to inhibit processing by Drosha/Pasha (Stefani et al., 2015); on the other hand, LIN-28 also promotes *SL1-LCE* trans-splicing, which results in down regulation of *pri-let-7* levels. LIN-28 could be either directly regulating LCE trans-splicing through its binding to *pri-let-7* or indirectly through other means such as regulating splicing components. In fact, in mammalian cells, LIN28 has been shown to indirectly affect alternative splicing by regulating the expression of certain splicing factors (Wilbert et al., 2012).

*lin-28(0)* animals exhibit greater than 100-fold elevation of *let-7* in the L1 and L2 stages, presumably as a result of a release of repression from both trans-splicing and inhibited biogenesis. When we reduced LCE trans-splicing by mutating SA1-6 in a wild-type *lin-28* background, mature *let-7* was not de-repressed as much as in *lin-28(0)*; although we observed a marked elevation in *pri-let-7,* consistent with a role for LCE trans-splicing in destabilizing *pri-let-7*, the attendant elevation of mature *let-7* was much more modest (only about 2-fold), apparently reflecting a potent inhibition by LIN-28 of *pri-let-7* Drosha/Pasha processing.

Interestingly, although we observed an elevation of mature *let-7* levels in *lin-28(0)* larvae that exceeded 100-fold, we observed no corresponding decrease in *pri-let-7*. Apparently, in the absence of LIN-28, the destabilization of *pri-let-7* levels on account of increased Drosha/Pasha processing is balanced by a commensurate stabilization of *pri-let-7* on account of reduced *SL1-LCE* trans-splicing. These observations further suggest that the level of *pri-let-7* in wild-type larvae could be subject to homeostatic regulation.

In mammals, trans-splicing is relatively rare compared to nematodes (Lei et al., 2016). However most human microRNA genes, including 10 of the 12 genes encoding *let-7fam* members, are located within introns of mRNAs or non-coding RNAs (Rodriguez et al., 2004, Kim and Kim, 2007). Therefore, it is possible that the spliceosomal machinery could contribute to the regulation of microRNA biogenesis in contexts other than *C. elegans let-7*. In fact, interplay between microRNA microprocessing and splicing has been previously observed. For example, the microRNA processing machinery Drosha/DGCR8, as well as microRNA primary transcripts, have been observed to be associated with supraspliceosomes, and inhibition of splicing can result in the elevation of microRNAs, including *let-7* (Agranat-Tamir et al., 2014). On the other hand, there is evidence of situations in which splicing and the biogenesis of intronic microRNAs can co-occur without apparently influencing each other (Kim and Kim, 2007), suggesting that connectivity between microRNA processing and host gene splicing is likely to be subject to regulation, depending on context and circumstances.

A previous study demonstrated that the *C. elegans* microRNA Argonaut ALG-1 could bind *in vivo* to the *let-7* locus LCE, suggesting that *let-7* miRISC could associate with *pri-let-7* in the nucleus and regulate *let-7* biogenesis (Zisoulis et al., 2012). In support of this idea, it was found that a transgene containing a mutant *let-7* locus that deleted 178 bp spanning the LCE displayed a marked decrease in *let-7* biogenesis compared to wild-type. When we generated the same 178 bp deletion in the endogenous *let-7* locus using genome editing, we also observed that the deletion resulted in decreased *let-7* expression, confirming that positive regulatory elements are contained in the 178 bp region. However, when we removed only the LCE at the endogenous locus, we observed no difference in *let-7* levels, indicating that the putative positive elements contained within the 178 bp deleted region are located outside of the LCE, and that the LCE itself does not exert a detectable positive role in *let-7* biogenesis. Our finding that LCE sequences are contained in a cytoplasmic mRNA that is produced by trans-splicing from *pri-let-7* has suggested that the LCE likely functions primarily by associating with *let-7fam* microRNAs in the cytoplasm. However, we cannot rule out the possibility that LCE sequences could also interact with miRISC in the nucleus.

Our results show that LIN-28-dependent trans-splicing of the *C. elegans let-7* primary transcript can act in *cis* to negatively influence *let-7* biogenesis, and at the same time produce a trans-acting inhibitory RNA, *SL1-LCE,* which negatively regulates the activity of *let-7fam* microRNAs. We propose that *SL1-LCE* functions as a sponge for *let-7fam* microRNAs through imperfect base pairing. Interestingly, animals with deletions of the LCE or splice acceptors displayed no overt phenotypes except in the sensitized background of *mir-48(0).* This suggests a that regulation of *SL1-LCE* production is not critical for normal development under standard laboratory conditions, but may function to modulate *let-7* biogenesis and *let-7fam* activity in the context of ensuring robust developmental timing under stressful physiological or environmental conditions.

## EXPERIMENTAL PROCEDURES

### Nematode methods and phenotypic analysis

*C. elegans* were cultured on nematode growth media (NGM) (Brenner 1974) and fed with *E. coli* HB101. Synchronized populations of developmentally staged worms were obtained by standard methods (Stiernagle 2006). Unless otherwise noted, all experiments were performed at 20°C.

For heterochronic phenotype analysis, early L4 animals were picked from healthy uncrowded cultures, placed onto individual plates seeded with HB101, and observed periodically until the end of the experiment. Fluorescence microscopy was used to score col-19::GFP expression.

### Sequence alignments, target prediction, and statistical significance

DNA and ORF alignments were performed using Clustal Omega (www.ebi.ac.uk).

*let-7fam*:LCE target predictions were performed using RNAhybrid (https://bibiserv2.cebitec.uni-bielefeld.de).

### Brood size counts

Young adult hermaphrodites were placed individually on plates seeded with HB101 and each animal was transferred daily to a fresh plate and the number of progeny produced on each plate was assessed until the animal stopped producing progeny.

### RNA extraction

Animals were collected and flash-frozen in liquid nitrogen, and total RNA was extracted using QIAzol reagent (Qiagen).

### Northern Blotting

Northern blotting was adapted from Lee and Ambros, 2001. RNA samples were run on 5% Urea-PAGE gels and then transferred to a GeneScreen Plus Hybridization membranes (PerkinElmer) by electrophoresis. After transfer, the membranes were cross-linked with 120mjoules of UV (wavelength of 254 nm) and baked at 80°C for one hour. Oligonucleotide probes were labeled using Integrated DNA Technologies’ (IDT) Starfire Oligos Kit with alpha-^32^P ATP and hybridized to the membranes at 37°C in 7% SDS, 0.2M Na2PO4 pH7.0 overnight. Membranes were washed at 37°C, twice with 2x SSPE, 0.1% SDS and twice with 0.5x SSPE, 0.1% SDS. The blots were exposed on a phosphorimager screen and imaged with a Typhoon FLA7000.

### Non-quantitative PCR

Samples of total RNA were pre-treated with turbo DNase (Invitrogen) (Pinto and Lindblad, 2010). cDNA was synthesized using SuperScript IV (Invitrogen) following the manufacturer’s instructions, with using the RT oligos “let-7 RT” or “oligo (dT)”. PCR was then performed using standard methods, and the products were analyzed by electrophoresis on a 2% agarose gel, imaged, cut out and purified from the gel, TA cloned (Invitrogen) and subjected to Sanger sequencing.

### Quantitative PCR

cDNA was synthesized as described above using the RT oligos “let-7 RT” and “gpd-1 QPCR R” (for full length *let-7* locus transcripts) or “pri-let-7 R” and “gpd-1 QPCR R” (for outron detection). qPCR reactions were performed using Qiagen QuantiFast SYBR Green PCR kit following the manufacturer’s instructions, using an ABI 7900HT Real Time PCR System (Applied Biosystems). With the exception of the experiments reported in Figure 6, Δ CTs were calculated by normalizing samples to *gpd-1* (*GAPDH*). Δ CTs were then inverted so that greater values reflect greater RNA levels, and were normalized to set the value of the least abundant sample to one.

### 5’ RACE

5’ RACE was adapted from (Pinto and Lindblad, 2010) and (Turchinovich et al., 2014). Samples of total RNA from late L2 animals were pre-treated with Turbo DNase (Invitrogen) (Pinto and Lindblad, 2010). Then 1.6 µl of the RNA (in H_2_O) was combined with 0.5 ul of 10 µM let-7 RT oligo, and 0.4 µl of 25 mM dNTPs. This mixture was incubated at 65°C for 10 minutes, chilled on ice, and then combined with 1.6 µl 25 mM MgCl_2_, 0.6 µl 100 mM MnCl_2_, 4 µl 5x First-Strand Buffer (Invitrogen), 2 µl 0.1 M DTT, 0.3 µl Ribolock and 8 µl H_2_O, and incubated at 42°C for two minutes, then 1.0 µl of SuperScript II (Invitrogen) was added and the mixture was incubated at 42°C for 30 minutes. 2.0 µl of TSO 5’RACE oligo (10 µM) was then added and the incubation was continued at 42°C for an additional 60 minutes. The reaction was heat inactivated at 70°C for 15 minutes, then diluted 1:10 and used for a standard PCR with the primers Rd1 SP and pri-let-7 R for *pri-let-7*, and Rd1 SP and LCE R for *SL1-LCE*. PCR products were TA-cloned (Invitrogen) and subjected to Sanger sequencing.

### Quantitative microRNA detection

microRNAs were quantified using FirePlex miRNA assay (Abcam) following the manufacturer’s instructions. Guava easyCyte 8HT (Millipore) was used for analysis. With the exception of the Calibrated RNA quantitation experiments (below), signals (arbitrary units) were normalized using geNorm (Vandesompele et al., 2002).

### Calibrated RNA quantitation

To generate T7 templates for the production of RNA standards corresponding to *pri-let-7* and *SL1-LCE*, the corresponding genomic sequences were PCR amplified from genomic DNA using the oligos T7 pri-let-7 F and let-7 RT, and T7 SL1-LCE F and let-7 RT, respectfully. T7 pri-let-7 added the T7 promoter to the *pri-let-7* PCR product, and T7 SL1-LCE added the T7 and SL1 sequences to the *SL1-LCE* PCR product. RNA from the respective PCR products was *in vitro* transcribed (IVT) following the manufacturer’s instructions (NEB’s HiScribe T7 High Yield RNA Synthesis Kit), and column purified. RNA concentration and quality was measured using an Advanced Analytics Fragment Analyzer. Known amounts of the IVT RNA were then serial diluted. cDNAs from the IVT serial dilutions, and from biological samples, were synthesized as described above and subjected to qPCR. Equal amounts of total RNA were used for each biological sample, and the amounts of *pri-let-7* and *SL1-LCE* in each biological sample were calculated from the standard curve generated from the IVT dilutions.

Synthetic oligonucleotides of *let-7, mir-48, mir-84,* and *mir-241* were ordered from IDT. Known amounts of these RNA oligonucleotides were serial diluted and subjected to FirePlex miRNA analysis along with biological samples. Equal amounts of total RNA were used for each biological sample, and the amounts each microRNA in each biological sample were calculated from the standard curve generated from the synthetic microRNA dilutions.

### Microscopy

Epifluorescence images were obtained with a Zeiss Imager.Z1 using the 10x objective.

### Targeted genome editing by CRISPR/Cas9

Mutants were generated using CRISPR/Cas9 methods adapted from Paix et al., 2014 and 2015. The germlines of young adult hermaphrodites were injected with a mix of CRISPR RNA (crRNA) targeting the region of interest in the *let-7* locus and the “co-CRISPR” marker *dpy-10,* tracrRNA, a single-stranded oligonucleotide HR template, Cas9 protein prepared as described in Paix et al., 2015, and water. F1 animals exhibiting the co-CRISPR phenotype were picked, allowed to lay eggs, and then genotyped by PCR. F2s were cloned from F1s that scored positively by PCR genotyping for the desired *let-7* locus modification. Homozygous F2s were then selected by PCR genotyping and subjected to Sanger sequencing for validation. All mutants were backcrossed to wild-type at least thrice. A list of the alleles generated along with the crRNAs used to generate them is below. Some crRNAs were generated using an IVT technique designed for this manuscript. crRNAs that were generated by IVT are noted in their names.

### *In vitro* transcription of crRNAs

To produce templates for the production of crRNAs by T7 in vitro transcription, equal amounts of two DNA oligonucleotides (100 µM) were mixed together: The sequence of the first oligonucleotide was the reverse compliment to the crRNA of interest followed by the reverse compliment of the T7 promoter; the second oligonucleotide (oCN183) was complimentary to the T7 promoter sequence of the first oligo. The oligonucleotide mixture annealed by rapidly heating to 95°C followed by cooling to 15°C over 10 minutes. 1.0 µl of the annealed oligonucleotide mixture was then added to an IVT reaction, and transcription carried out following the manufacturer’s instructions (NEB). The RNA product was then column purified. **CAAAACAGCATAGCTCTAAAACNNNNNNNNNNNNNNNNNNNCCTATAGTGAGTC GTATTA reverse compliment of universal sequence 19 nt reverse compliment of guide reverse compliment of T7 promoter**

### Summary of CRISPR/Cas9 alleles

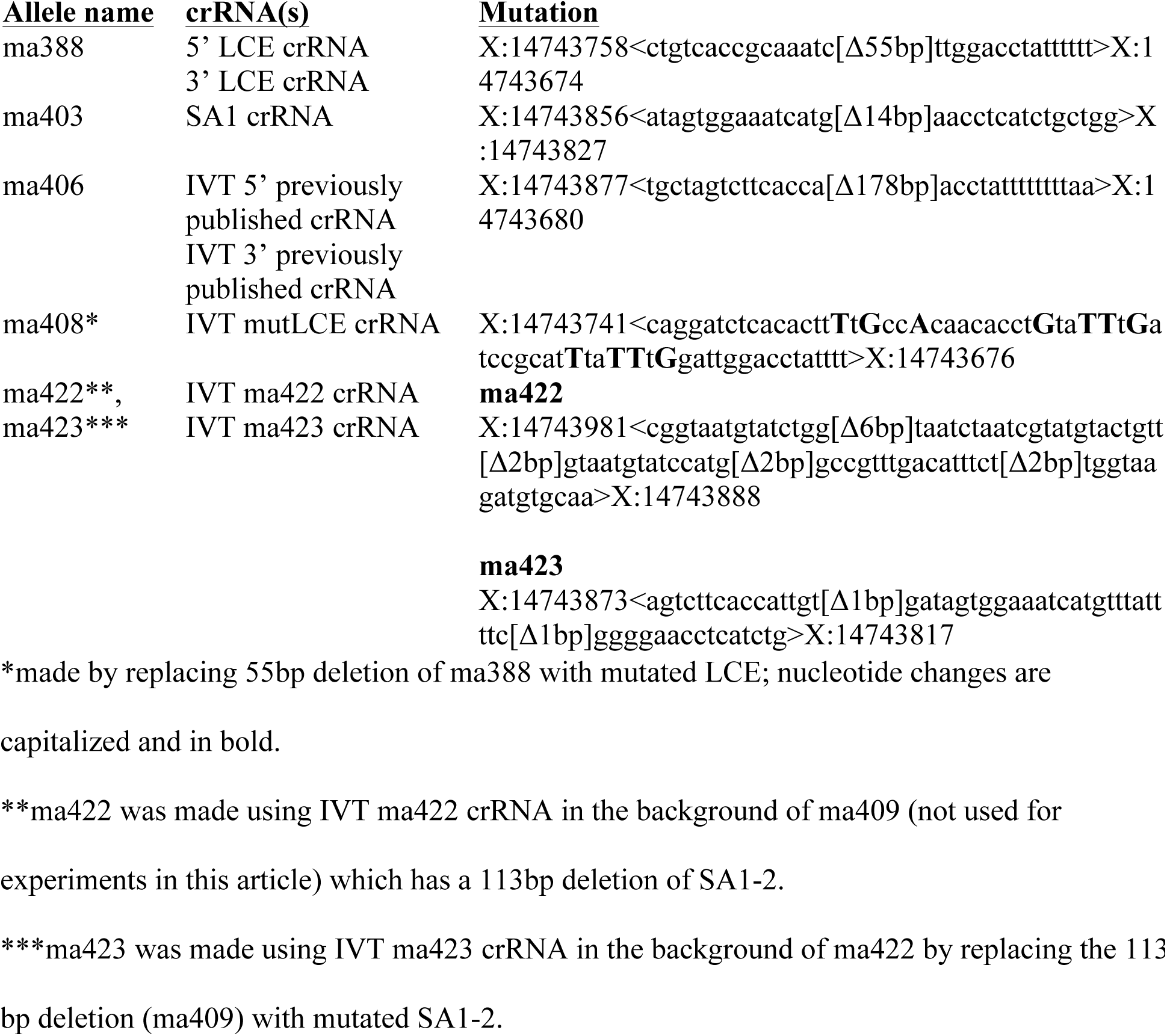

### Transgenic construct

The pCN30 construct, which contains the *let-7* locus’ ORF tagged with GFP on its C-terminus, was constructed by cloning GFP into the *let-7* genomic rescue plasmid pZR001 (Ren and Ambros, 2015).

### Oligos

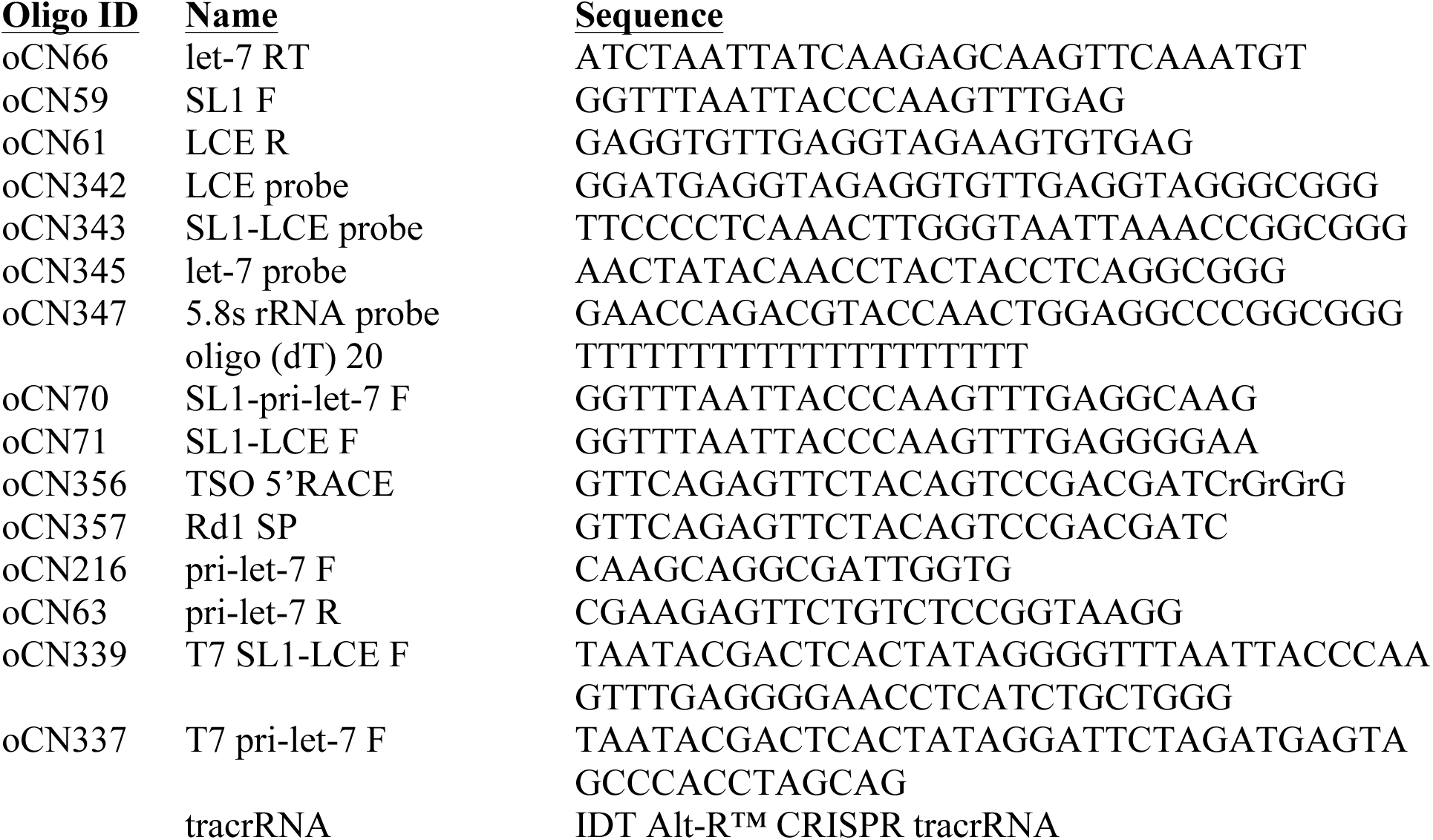
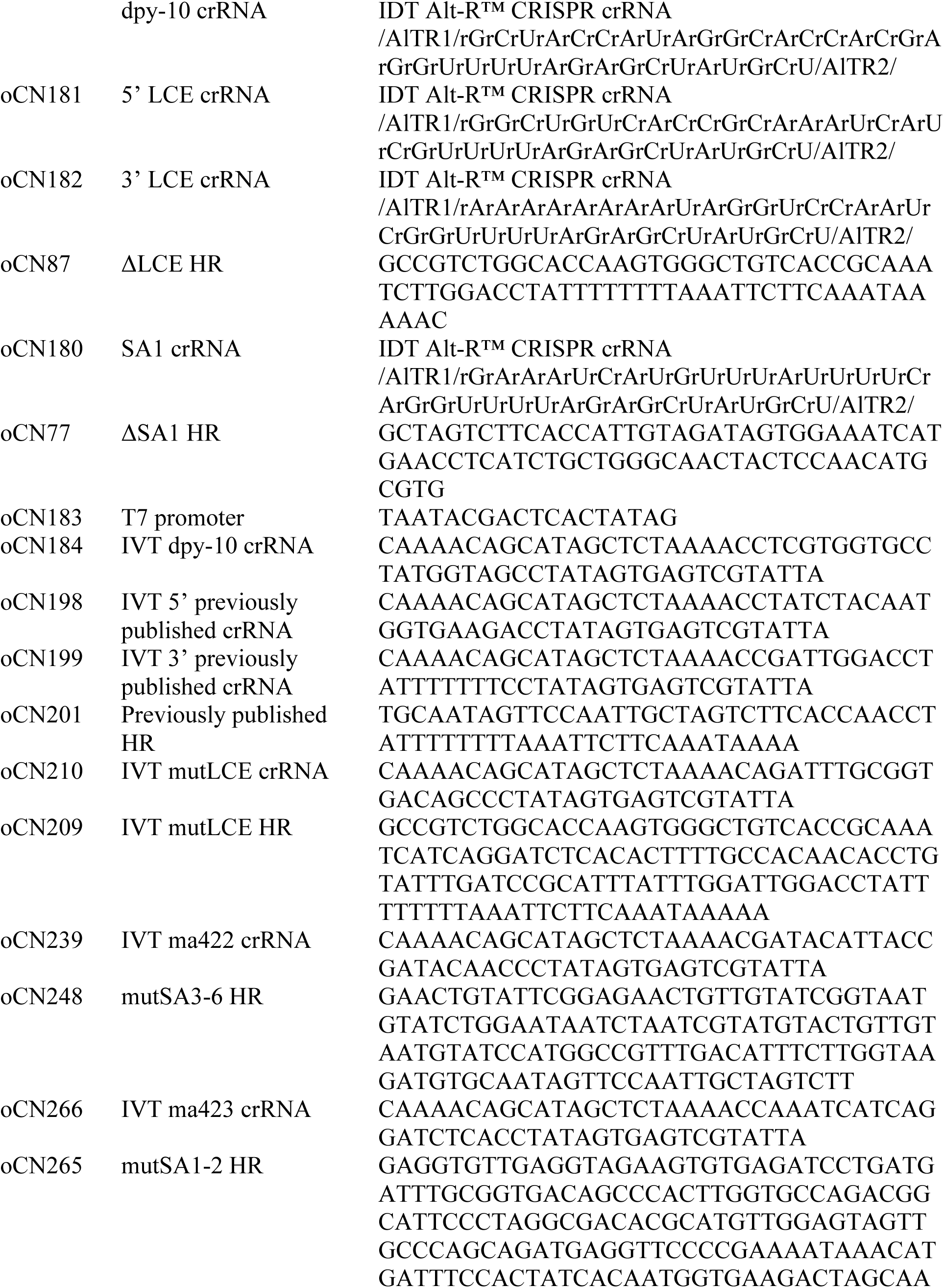
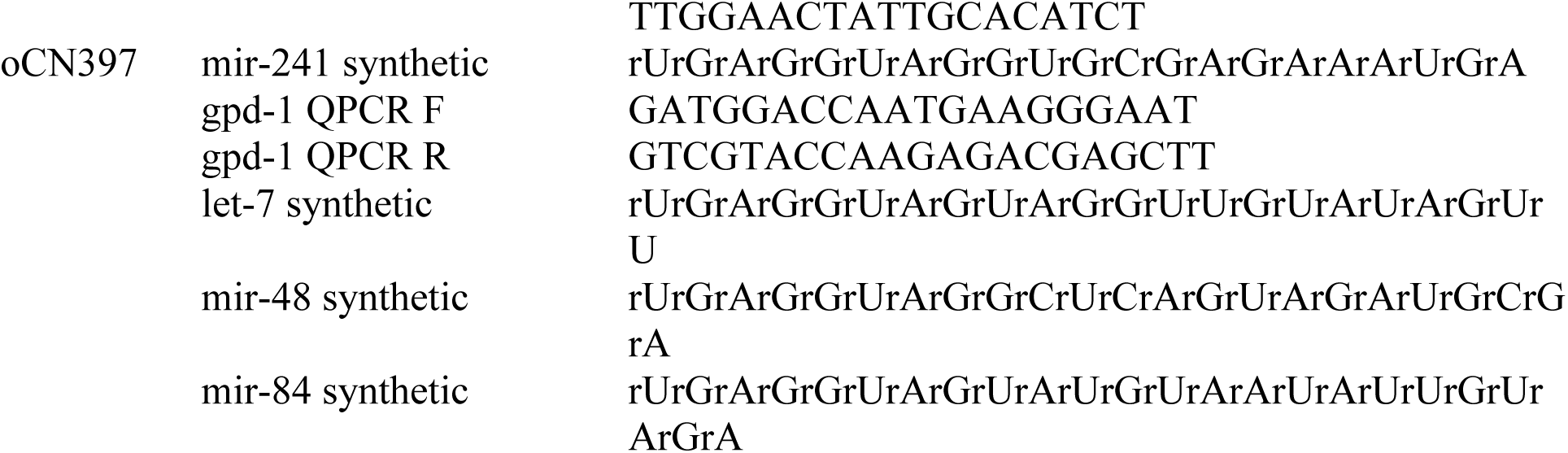

### *C. elegans* strains

***C. elegans* strains** **Strain**
| **<u>Name</u>** | **<u>Strain Description</u>** | **<u>Genotype</u>** |
| --- | --- | --- |
| MT355 | *lin-14(gf)* | *lin-14(n355)* X |
| VT965 | *lin-14(lf)* | *lin-14(n179)* X |
| VT1295 | *lin-28(0)/lin-28* null | *lin-28(n719)* I; *maIs105* V |
| VT1367 | wild-type/WT | *maIs105* [*col-19::gfp*] V |
| VT3594 | *lin-28(0); lin-46(0)* | *lin-28(n719)* I; *lin-46(maIs105)* V |
| VT3609 | *lin-4(lf)* | *lin-4(e912)* II; *maIs105* V |
| VT3616 | ΔLCE | *let-7(ma388)* X; *maIs105* V |
| VT3666 | ΔSA1 | *let-7(ma403)* X; *maIs105* V |
| VT3669 | previously published deletion | *let-7(ma406)* X; *maIs105* V |
| VT3675 | ΔSA1; *lin-28(0)* | *let-7(ma403)* X; *lin-28(n719)* I; *maIs105* V |
| VT3678 | mutLCE | *let-7(ma408)* X; *maIs105* V |
| VT3718 | mutSA1-6 | *let-7(ma422, ma423)* X; *maIs105* V |
| VT3719 | mutLCE; *mir-48(0)* | *let-7(ma408)* X; *miR-48(n4097)*, *maIs105* V |
| VT3720 | ΔLCE; *mir-48(0)* | *let-7(ma388)* X; *miR-48(n4097)*, *maIs105* V |
| VT3721 | ΔLCE; *mir-48(0)*, *mir-241(0)* | *let-7(ma388)* X; *miR-48*, *miR-241(nDF51)*,<br>*maIs105* V |
| VT3837 | *mir-48(0)* | *mir-48(n4097)*, *maIs105* V |
| VT3838 | mutSA1-6; *mir-48(0)* | *let-7(ma422, ma423)* X; *mir-48(n4097)*, *maIs105* V |
| VT3839 | *let-7* locus' ORF::GFP<br>transgene | *maEx264* [*unc-119+*; *plet-7::let-7* locus with<br>*ORF::GFP*]; *unc-119(ed3)* III |
| VT3840 | *mir-48(0)*, *mir-241(0)* | *mir-48*, *mir-241 (nDf51)*, *maIs105* V |

## ACKNOWLEDGEMENTS

We thank the members of the Ambros and the Mello laboratories for helpful discussions. We also thanks the Mello lab for help with the northern blotting, especially Takao Ishidate. This work was supported in part by a Translational Cancer Biology Training Grant, NIH T32 training grant # T32CA130807-06A1, and by NIH R01GM34028.

## AUTHOR CONTRIBUTIONS

C.N. performed the experiments and analyzed the data. C.N. and V.A. designed the experiments and wrote the paper.

